# Expanded protocadherin-1 usage reveals a broader hantavirus entry landscape

**DOI:** 10.64898/2026.06.23.734139

**Authors:** Cierra Word, Nahomi Guerra-Pilaquinga, Ezgi Kasikci, Lohit Khera, Ramandeep Kaur, Upendra P. Lambe, Maria Eugenia Dieterle, Kartik Chandran, Rohit K. Jangra

**Affiliations:** Department of Microbiology and Immunology, Center for Applied Immunology and Pathological Processes, Center for Cardiovascular Diseases and Sciences, Louisiana State University Health Sciences Center-Shreveport, Shreveport, Louisiana, USA; Department of Microbiology and Immunology, Albert Einstein College of Medicine, Bronx, New York, USA

**Author notes:** Corresponding author: Rohit K Jangra.

## Abstract

Mammalian hantaviruses are RNA viruses that cause hantavirus cardiopulmonary syndrome in the Americas and hemorrhagic fever with renal syndrome in Eurasia. The cellular entry mechanisms of most hantaviruses remain poorly defined. To examine entry by phylogenetically distinct hantaviruses, we generated replication-competent recombinant vesicular stomatitis viruses (rVSVs) bearing Gn/Gc proteins from Necoclí, Sangassou, Thottapalayam, Kenkeme, Nova, Oxbow, and Tula viruses. All these Gn/Gc proteins except Kenkeme supported infection of primary human endothelial cells, indicating that endothelial cell entry is permissive for a broader range of hantaviruses than previously appreciated. Except for rVSV-Kenkeme, these rVSVs did not acquire additional mutations beyond pre-engineered rescue-enhancing changes during rescue and passaging. Genetic studies in human cells lacking protocadherin-1 (PCDH1) showed that Necoclí, Tula, and Nova viruses use PCDH1 for efficient infection, although the Nova phenotype was weaker. These three Gn/Gc proteins bound soluble PCDH1 with different apparent avidities, and infection by the corresponding rVSVs was inhibited by soluble PCDH1; Necoclí and Tula, but not Nova, were also blocked by a PCDH1-targeting monoclonal antibody. Authentic Tula virus infection was similarly reduced in PCDH1 knockout endothelial cells. Finally, the broadly reactive anti-Gn/Gc human monoclonal antibody ADI-42898 efficiently neutralized Necoclí, Nova, Sangassou, and Kenkeme rVSVs but showed weak or undetectable activity against Oxbow, Tula, and Thottapalayam rVSVs. Together, these findings expand the range of hantavirus glycoproteins capable of mediating infection of human endothelial cells, broaden the phylogenetic scope of PCDH1-dependent entry, and identify receptor-targeted and viral glycoprotein-targeted strategies with differential activity.

**Importance:** Many newly discovered hantaviruses are known only from sequence data, leaving their ability to enter human cells and their receptor usage unresolved. Using a BSL2-compatible rVSV system, we show that glycoproteins from several divergent hantaviruses can mediate infection of primary human endothelial cells, indicating that endothelial cell entry is permissive for a broader range of hantaviruses than previously appreciated. We also show that protocadherin-1 (PCDH1), previously linked mainly to New World hantaviruses, is used by Necoclí, Tula, and Nova viruses but not universally across the panel, revealing broader but heterogeneous receptor usage. An authentic Tula virus experiment supports this conclusion beyond the surrogate system. Finally, a broadly reactive anti-Gn/Gc antibody neutralizes several, but not all, of these viruses, highlighting both the promise and the limits of broadly protective countermeasures and the utility of these rVSVs for evaluating entry inhibitors.

## Introduction

Hantaviruses are enveloped, negative-sense RNA viruses that belong to the family *Hantaviridae* in the order *Elliovirales*(1). When humans are accidentally exposed to aerosolized excreta from infected reservoir hosts, rodent-borne hantaviruses can cause severe, often fatal diseases, such as hantavirus cardiopulmonary syndrome (HCPS) in the Americas and hemorrhagic fever with renal syndrome (HFRS) in Europe and Asia(2, 3). Periodic hantavirus outbreaks continue to occur, and Andes virus (ANDV) is notable for rare human-to-human transmission in a close-contact setting, such as on the MV *Hondius* Cruise Ship in April-May 2026(4–7). Despite their significant impact on public health, there are currently no FDA-approved vaccines or therapeutics for hantavirus infections, and our understanding of their entry mechanisms remains poor(2, 8).

Studies based on genetic sequencing over the past few decades have discovered many new mammalian hantaviruses carried by rodents, insectivores, and bats(9, 10). Currently, the International Committee on Taxonomy of Viruses (ICTV) recognizes 48 species of mammalian hantaviruses globally(11). However, for most of these hantaviruses, no virus isolates are available, and their potential to cause human infection and illness remains unknown(12). Additionally, the need for high-level biocontainment (BSL3) in hantavirus research and the lack of a robust reverse genetics systems to study the complete hantavirus infection cycle greatly hinder our understanding of their biology(2).

Hantavirus particles carry a negative-sense, tripartite genome composed of small (S), medium (M), and large (L) segments(13). The S segment encodes the nucleoprotein (N), which coats the genome; the L segment encodes the L protein, which possesses RNA-dependent RNA polymerase and endoribonuclease activities required for genome replication and transcription, respectively. The M segment encodes the glycoprotein complex as a single polyprotein, which is processed by cellular proteases to generate the Gn and Gc glycoproteins(14). The Gn/Gc heterodimers are arranged in a tetrameric cloverleaf-like structure on hantavirus particles, forming spikes(15). Gn is believed to interact with host receptors, while Gc mediates fusion of the viral and endosomal membranes during cellular entry(2, 15). Because Gn/Gc are necessary and sufficient to mediate cellular entry, vesicular stomatitis viruses carrying Gn/Gc serve as powerful tools for studying viral entry(2, 16).

Among proposed hantavirus entry receptors, only protocadherin-1 (PCDH1) has compelling *in vivo* support; it is required for infection and pathogenesis of ANDV in Syrian hamsters(17, 18). Earlier *in vitro* studies have implicated β1-3 integrins (ITGβ1, ITGβ2, ITGβ3), decay-accelerating factor (DAF)/CD55, and the globular head of C1q receptor (gC1qR) as potential attachment factors or receptors for hantavirus entry(19–24). However, their *in vivo* relevance is not clear, and ITGβ1, ITGβ3 and DAF/CD55 are not essential for hantavirus infection in human endothelial cells, the major targets of hantavirus infection(25). Genetic knockout and biochemical studies demonstrated that protocadherin-1 (PCDH1) is required for the entry of 5 species of New World hantaviruses: Andes virus, Sin Nombre virus (SNV), Prospect Hill virus, Maporal virus, and Choclo virus, but not for the Old World hantaviruses such as Hantaan virus (HTNV), Puumala virus, and Seoul virus(18, 25). The first extracellular cadherin (EC1) domain of PCDH1 binds the Gn/Gc of New World hantaviruses, and disruption of the PCDH1:Gn/Gc interaction by complete knockout or point mutations in EC1 protects Syrian hamsters against a lethal Andes virus challenge(17, 18). Moreover, soluble PCDH1 carrying the EC1 domain and the EC1-specific monoclonal antibody (mAb-3305) neutralize rVSV-ANDV and SNV infections(17, 18). However, the extent of PCDH1 usage by distinct hantavirus lineages remains unknown.

To capture genetic, geographic, and reservoir-host variation, we focused on shrew-borne Thottapalayam virus (TPMV) and Kenkeme virus (KKMV); mole-borne Nova virus (NVAV), and Oxbow virus (OXBV); New World rodent-borne Necoclí virus (NECV); Old World rodent-borne Sangassou virus (SANGV), and vole-borne Tula virus (TULV)(26). Thottapalayam virus was first isolated from the Asian house shrew in India(27), and Kenkeme virus was identified in the flat-skulled shrew in Russia(28). Nova virus is a highly divergent mole-borne virus of the European mole and was initially detected in Hungary, with high prevalence later documented in France(29, 30). Necoclí virus is principally associated with Cherrie’s cane rat in Colombia(31). Sangassou virus is the first indigenous African hantavirus isolate and was identified in the African wood mouse in Guinea(32). Tula virus is classically associated with common voles and related arvicoline hosts across Eurasia(26, 33). Among these viruses, Sangassou virus entry studies implicated integrin β1 as a receptor(32), whereas Tula virus has the strongest direct evidence of human relevance, with a molecularly confirmed symptomatic infection reported in an immunocompetent patient(34). While serologic evidence suggests possible human exposure to Thottapalayam and Sangassou viruses, the ability of Kenkeme, Nova, Oxbow, and Necoclí viruses to cause human infection or disease remains unknown(35–38). Although their reservoir-host associations are better established, human-cell tropism and receptor usage remain poorly characterized for most of these viruses(26, 38). Together, these phylogenetically divergent hantaviruses associated with a variety of reservoir hosts provide a framework to evaluate human cell-tropism and PCDH1 dependence across mammalian hantaviruses.

Here, we generated rVSVs expressing the Gn/Gc proteins of the seven genetically and ecologically heterogeneous hantaviruses mentioned above to investigate their entry mechanisms and host receptor usage. Infectivity assays in human cells, including primary endothelial cells, demonstrated that these Gn/Gc, except that of Kenkeme virus, can mediate infection of human cells without acquiring additional adaptive mutations. Experiments with PCDH1 knockout human cells and ELISA-based binding assays demonstrated that Necoclí, Tula, and Nova virus Gn/Gc-mediated infections depend on PCDH1. Finally, we demonstrated the utility of PCDH1 and Gn/Gc-targeting approaches for neutralizing these viral infections.

## Results

### Divergent hantavirus Gn/Gc proteins mediate infection of human endothelial cells

Hantavirus Gn/Gc sequences are highly divergent across species, with 40-60% amino acid differences between them. Our hantavirus panel was selected to sample viruses spanning diverse reservoir hosts, phylogenetic lineages and geographic origins (Fig. 1). To enable cellular entry studies of these hantaviruses at BSL2, we generated recombinant vesicular stomatitis viruses (rVSVs) expressing Gn/Gc in place of the native VSV G protein. These viruses also express an enhanced GFP reporter (Fig. 2A, top panel).

**Fig. 1.**
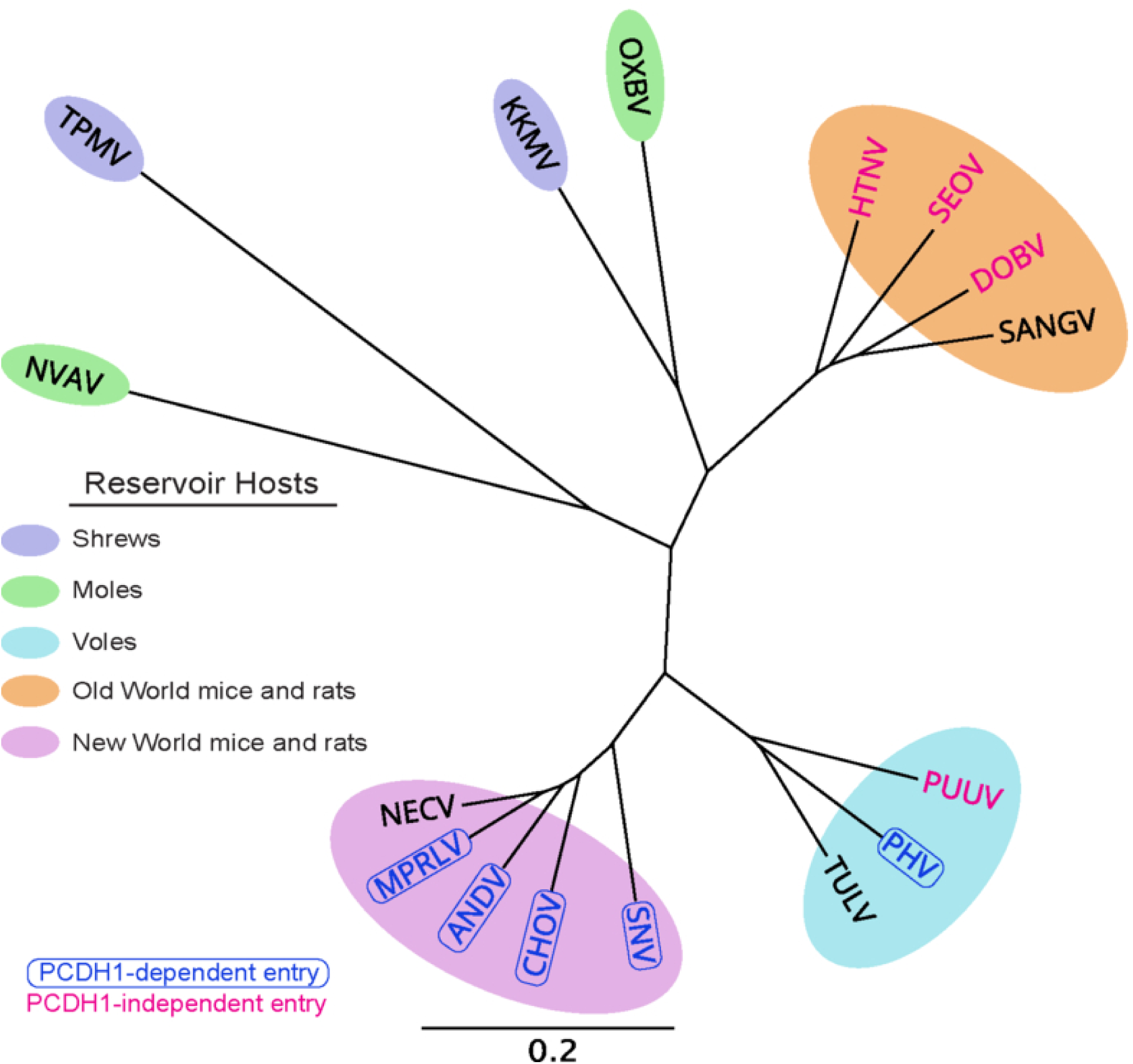
Phylogenetic relationships, reservoir hosts, and known PCDH1 usage of hantaviruses included in this study. Neighbor-joining phylogenetic analysis performed using the full-length amino acid sequences of the glycoprotein precursor (GPC) of the indicated hantaviruses in Geneious Prime v2025.2. Amino acid sequences were translated from the nucleotide sequences obtained from GenBank under the following accession numbers: Andes virus (ANDV, AF291703.2), Hantaan virus (HTNV, M14627.1), Maporal virus (MPRLV, AY363179.1), Sin Nombre virus (SNV, L37903.1), Choclo virus (CHOV, DQ285047.1), Prospect Hill virus (PHV, X55129.1), Puumala virus (PUUV, M29979.1), Seoul virus (SEOV, S47716.1), Dobrava-Belgrade virus (DOBV, AJ410616.1), Necoclí virus (NECV, KF494345.1) Tula virus (TULV, Z69993.1), Nova virus (NVAV, KR072622.1), Oxbow virus (OXBV, FJ539167.1), Kenkeme virus (KKMV, KJ857337.1), Thottapalayam virus (TPMV, DQ825771.1), and Sangassou virus (SANGV, JQ082301.1). Reservoir hosts of the included viruses are indicated by colors ovals. Viruses known to use PCDH1 are marked by blue open boxes and blue text, whereas viruses known not to use PCDH1 are marked in red.

**Fig. 2.**
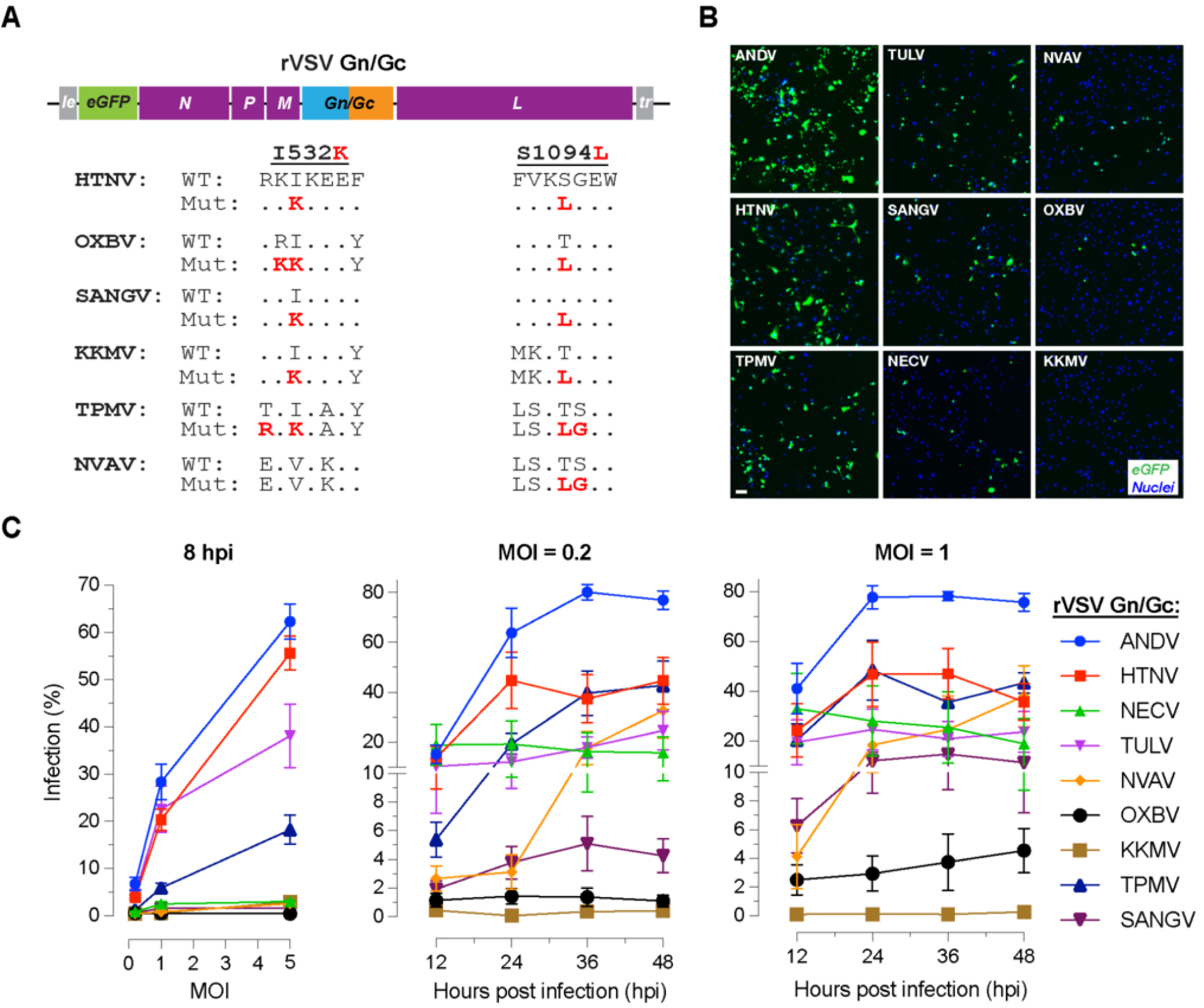
Divergent hantavirus Gn/Gc proteins mediated infection of human endothelial cells. **(A)** Top, schematic of the recombinant vesicular stomatitis virus (rVSV) genome, in which hantavirus Gn/Gc proteins replace the native VSV glycoprotein G. These viruses also encode an enhanced green fluorescent protein (eGFP) reporter in the first transcriptional unit. Bottom, alignment of Gn/Gc regions across hantaviruses, highlighting in bold the I532K substitution in the cytoplasmic tail of Gn and the S1094L substitution in the Gc ectodomain of Hantaan virus (HTNV) Gn/Gc, which we previously showed enhances rVSV recovery. For some Gn/Gc proteins, nearby residues were also changed to match the HTNV sequence. Wild-type (WT) sequences and pre-engineered amino acid substitutions (Mut) into the indicated Gn/Gc are shown in red. Dots indicate amino acids identical to those in HTNV. **(B)** Primary human pulmonary microvascular endothelial cells (HPMECs) infected with the indicated rVSVs at a multiplicity of infection (MOI) of 1 and fixed 24 hours later. Nuclei were counterstained with Hoechst-33342, and infection was assessed by eGFP expression. Representative images from one of four experiments are shown. Scale bar = 100 μm. **(C)** Left, HPMECs were infected with the indicated rVSVs at MOIs of 0.2, 1, or 5, fixed at 8 hours and imaged as in (B). Nuclei and infection were quantified using an automated Cytation 5 Cell Imaging Multi-Mode Reader (Agilent BioTek) with Gen5 software. Results are expressed as percent infection (n = 6-16) from 3-6 independent experiments. Middle and right, HPMECs were infected at MOIs of 0.2 and 1, respectively, and infection was measured at the indicated time points. For MOI 0.2, n = 6-8 from 4 independent experiments; for MOI 1, n = 4-6 from 3 independent experiments. ANDV, Andes virus; TULV, Tula virus; NECV, Necoclí virus; OXBV, Oxbow virus; SANGV, Sangassou virus; KKMV, Kenkeme virus; TPMV, Thottapalayam virus; NVAV, Nova virus. Each n represents one well of a 96-well plate, with 500-1,000 cells scored per condition. Data are shown in mean ± SEM; some error bars are smaller than the symbols.

Because New World hantavirus glycoproteins are more compatible with plasma-membrane budding, whereas Old World hantavirus glycoproteins are typically Golgi-localized, rVSVs bearing New World hantavirus Gn/Gc are generally rescued without adaptive mutations(39, 40). In contrast, Old World hantaviruses, such as Hantaan virus and Dobrava-Belgrade virus, require two mutations, I532K in the cytoplasmic tail of Gn, and S1094L in the membrane-proximal part of the Gc ectodomain, to enhance rVSV generation(16). These mutations are unlikely to directly affect receptor engagement because they were previously shown to enhance rVSV infectivity primarily by altering glycoprotein localization and incorporation into virions(16). Accordingly, we preemptively altered the Gn/Gc sequences of Thottapalayam, Kenkeme, Nova, Oxbow, and Sangassou viruses to incorporate HTNV-adaptive changes for rVSV generation (Fig. 2A, bottom panel).

Plasmids encoding cognate rVSV genomes carrying codon-optimized Gn/Gc sequences for expression in human cells were transfected into 293FT cells together with the helper plasmids, and the resulting supernatants were used to infect Huh7.51 cells for viral amplification and stock generation. These viruses reached titers of 5-6 log_10_ per mL in Huh7.5.1 cells. Sequencing of the Gn/Gc inserts in the rVSV genomes confirmed that none of the rVSVs, except rVSV-Kenkeme, acquired additional mutations beyond the pre-engineered ones. In rVSV-Kenkeme, a G485R mutation, numbered according to the glycoprotein precursor, was detected, although its significance remains unclear.

To compare infectivity, we normalized viral input by titrating the rVSVs on permissive Huh7.5.1 cells. These titers were then used to determine the multiplicity of infection (MOI) for infection of primary human pulmonary microvascular endothelial cells (HPMECs), a major target cell type for hantavirus infection. HPMECs were infected at an MOI of 1 and fixed 24 hours later to assess eGFP-positive cells. All rVSVs except rVSV-Kenkeme infected these cells, although with varying efficiency (Fig. 2B). For quantitative analysis, we evaluated infection at MOIs of 0.2, 1, and 5 at 8 hours post-infection. At an MOI of 0.2, only rVSV-Tula infection was evident (4.68%). At MOIs of 1 and 5, rVSV-Tula (23% and 38%, respectively) and rVSV-Thottapalayam (6% and 18%, respectively) infection was detectable, along with the positive controls rVSV-ANDV and rVSV-HTNV, whereas the remaining viruses were not (Fig. 2C, left panel).

Kinetic experiments over 48 hrs at MOIs of 0.2 and 1 (Fig. 2C, middle and right panels, respectively) further supported the higher infectivity of rVSV-Thottapalayam and rVSV-Tula, and also revealed substantial infection by rVSV-Necoclí (18 and 36% at 12 hours, respectively) and rVSV-Nova (32 and 38% at 48 hours, respectively). The other three rVSVs, rVSV-Sangassou, rVSV-Oxbow, and rVSV-Kenkeme, showed much lower infectivity: rVSV-Sangassou peaked at 11%, rVSV-Oxbow at 5%, and rVSV-Kenkeme remained barely detectable (<2%). Three broad kinetic patterns were observed: rVSV-Thottapalayam, rVSV-Nova, and, to a lesser extent, rVSV-Sangassou showed a steady increase over time, similar to rVSV-ANDV and rVSV-HTNV; rVSV-Necoclí and rVSV-Tula showed relatively higher infection with little increase over time; and rVSV-Kekeme and rVSV-Oxbow showed very low infection with no increase over time (Fig. 2C). Together, these results show that most of these divergent hantavirus Gn/Gc proteins can mediate infection of primary human endothelial cells. Except Kenkeme virus, this activity was retained without acquisition of additional mutations beyond the pre-engineered changes during rVSV generation.

### PCDH1 knockout reduces infection by rVSVs bearing Necoclí, Tula, and Nova virus Gn/Gc-mediated entry

We have previously used PCDH1 knockout human osteosarcoma U2OS cells to evaluate PCDH1’s role in hantavirus infection(17, 18). To assess PCDH1 usage by these rVSVs, we infected WT U2OS cells, a clonal PCDH1 knockout (KO) U2OS cell line, and PCDH1 KO U2OS cells complemented with PCDH1 cDNA with pre-titrated amounts of rVSVs selected to yield 20-30% infection in WT cells, except for rVSV-Kenkeme, for which the target infection level was 10%, and quantified infection 24 hours later (Fig. 3A). As with rVSV-ANDV, our positive control, rVSV-Tula infection was reduced in PCDH1 KO cells, and complementation restored infection. Although rVSV-Necoclí and rVSV-Nova infection was also reduced in PCDH1 KO cells, it was not restored in the complemented U2OS cells. Infection by the other 4 rVSVs was unaffected by PCDH1 status.

**Fig. 3.**
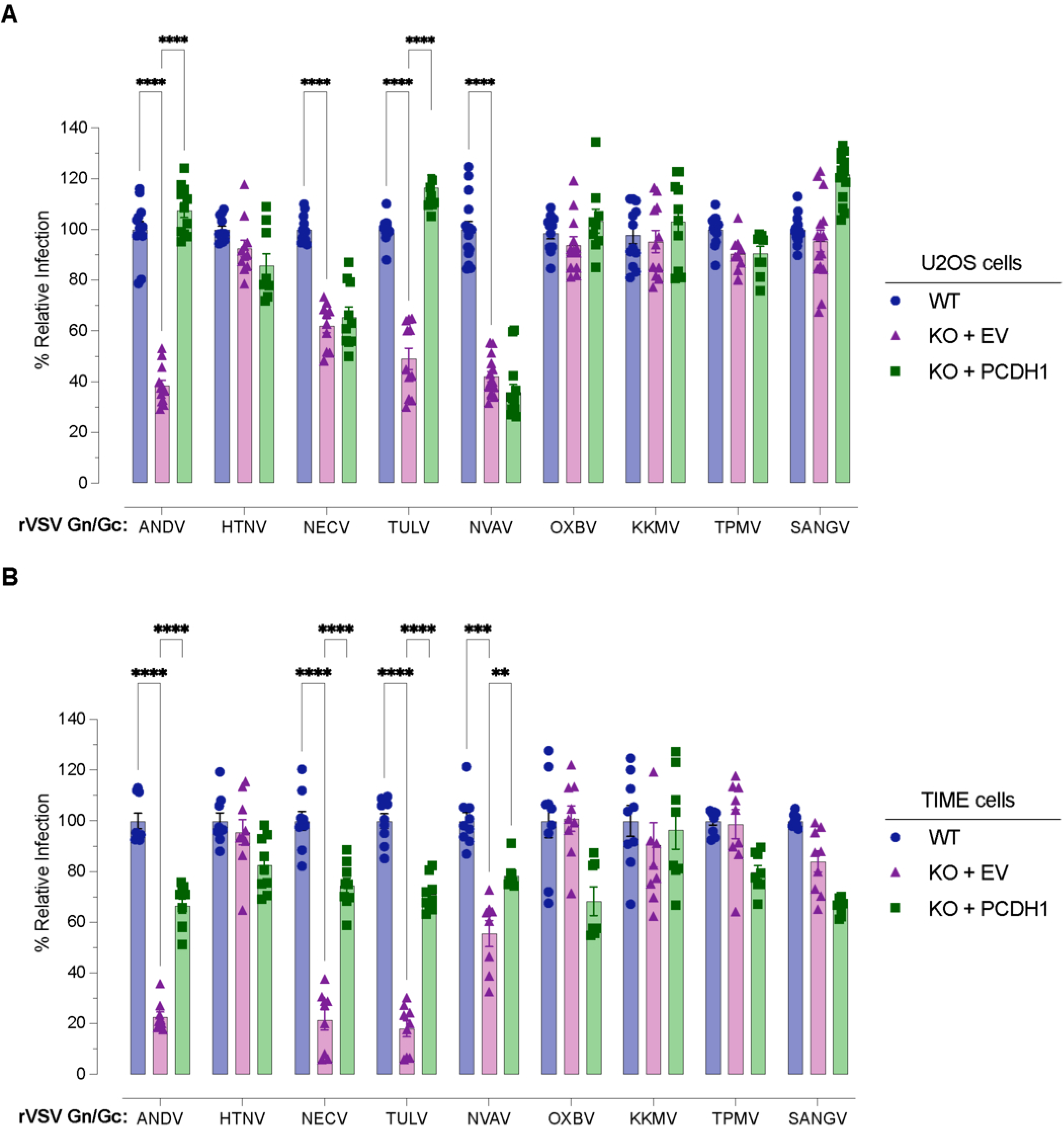
PCDH1 knockout reduces infection of rVSVs bearing Necoclí, Tula, and Nova glycoproteins. **(A)** Wild-type (WT) U2OS cells, a clonal PCDH1 knockout (KO) cell line carrying empty vector (KO + EV), and PCDH1 KO U2OS cells complemented with PCDH1 cDNA (KO + PCDH1) were infected with the indicated rVSV and infection was quantified 24 hours later as in Fig. 2C. **(B)** WT TIME cells, a mixed PCDH1 KO TIME cell population carrying empty vector (KO + EV), and PCDH1 KO TIME cells complemented with PCDH1 cDNA (KO + PCDH1) were infected with the indicated rVSV and infection was quantified 12 hours later as in Fig. 2C. Viruses were pre-titrated to estimate inocula for each rVSVs that yielded 20-30% infection in WT U2OS and TIME cells. except for rVSV-Kenkeme, for which the target infection level was approximately 10%. Data are presented as relative infection, with WT set to 100%. Each dot represents infection measured from 1,000-2,000 U2OS cells or 500-1,000 TIME cells per condition. Data are shown as mean ± SEM (n = 9-16 from 3-6 independent experiments for U2OS cells; n = 7-9 from 3 independent experiments for TIME cells). Statistical comparisons were performed using Two-Way ANOVA with a mixed-effects model and the Geisser-Greenhouse correction. ****, p<0.0001; ***, p=0.0009; **, P=0.0027. Nonsignificant differences are not indicated. Some error bars are smaller than the symbols.

To verify these results in endothelial cells, we tested the same rVSV infections in human TIME endothelial cells, as in our previous work(25). The results in TIME cells were largely consistent with those in U2OS cells: rVSV-ANDV infection decreased by about 80%; rVSV-Necoclí and rVSV-Tula infection each decreased by about 80%; and rVSV-Nova infection decreased by about 50% in the absence of PCDH1 (Fig. 3B). Importantly, infection of all 4 viruses was restored by PCDH1 complementation in this nonclonal knockout population. In contrast, rVSV-Oxbow, rVSV-Kenkeme, rVSV-Thottapalayam, and rVSV-Sangassou infections were unaffected by PCDH1. These results demonstrate that PCDH1 ablation in human cells reduces infection mediated by Necoclí, Tula, and Nova Gn/Gc.

### Necoclí, Tula, and Nova Gn/Gc specifically bind to PCDH1

To determine whether these Gn/Gc bind PCDH1, we first normalized input rVSVs by measuring the VSV matrix (M) protein by Western blotting (Fig. 3A, top panel). We then used our previously validated ELISA, in which rVSV particles bearing hantavirus Gn/Gc are captured by the soluble EC1-2 (sEC1-2) domain of PCDH1 immobilized on an ELISA plate (17, 18). As with rVSV-ANDV, binding of rVSV-Necoclí to PCDH1 was dose dependent, although of lower avidity (Fig. 4A, bottom panel). Binding of rVSV-Tula and rVSV-Nova was weaker and was detectable only at higher virion inputs (Fig. 4B).

**Fig. 4.**
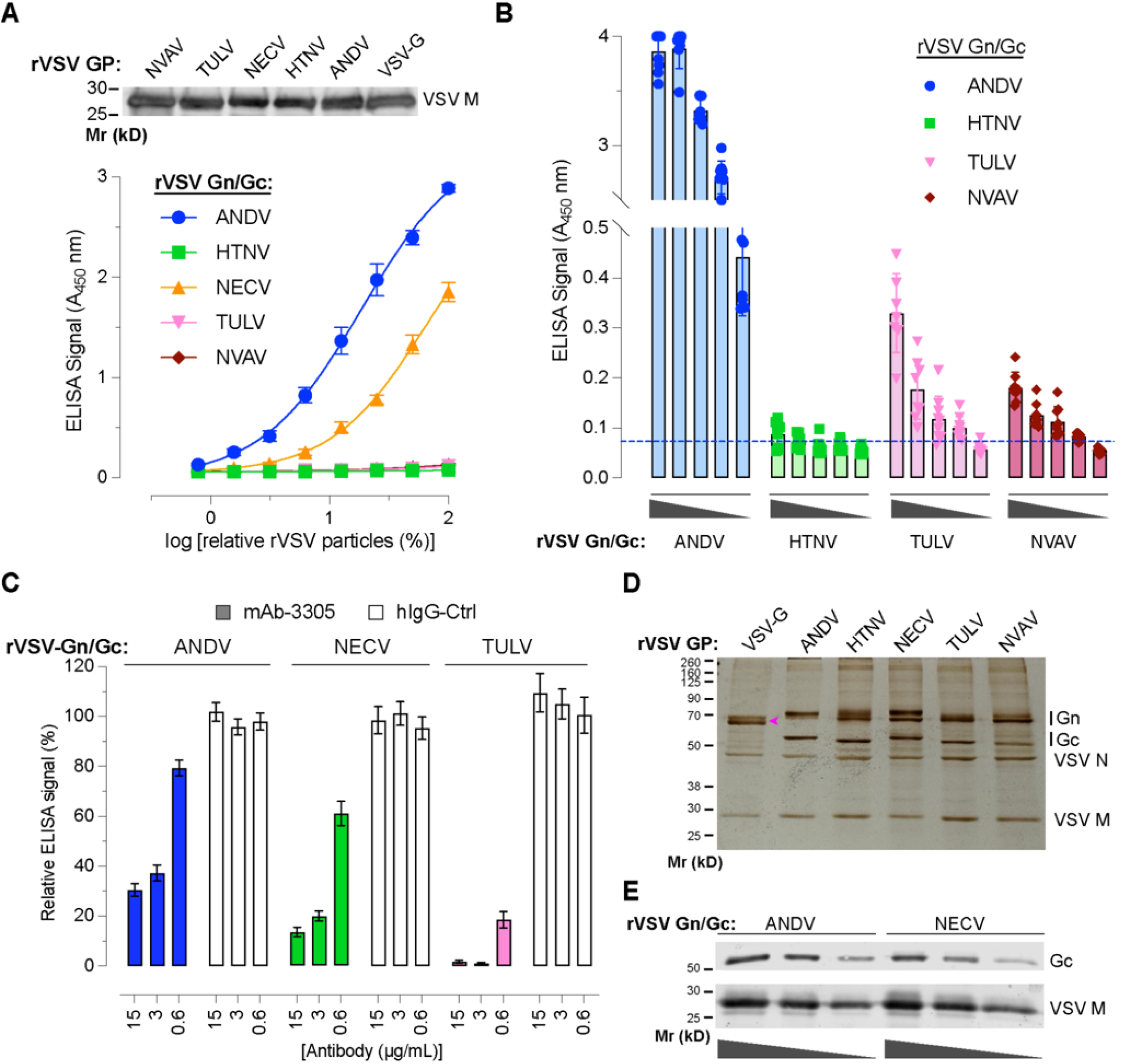
Necoclí, Tula, and Nova Gn/Gc specifically bind to PCDH1. **(A)** Top, a representative Western blot showing normalized amounts of the indicated pelleted rVSV particles, estimated by VSV matrix (M) protein levels, used as input for the sEC1-2 capture ELISA. Bottom, normalized amounts of FSL-biotin-labeled rVSVs were applied to sEC1-2-coated plates, and bound virus was quantified by detection of biotin embedded in the viral membrane using a streptavidin-HRP-based ELISA. Data are shown as mean ± SEM (n = 11-12) from 4 independent experiments. **(B)** Bar graph of the sEC1-2-capture ELISA, as in (A), except that higher amounts of rVSVs were tested. Data are shown as mean ± SEM (n = 7-9) from 3 independent experiments. The dotted blue line indicates the cutoff for a positive signal, defined as the mean + 3 standard deviations of the ELISA signal obtained with the lowest concentration of the negative control, rVSV-HTNV, tested. **(C)** EC1-specific mAb-3305 or unrelated hIgG-Ctrl at the indicated concentrations was preincubated with sEC1-2-coated plates before binding and detection of the indicated FSL-biotin-labeled rVSVs by ELISA. ELISA signal was normalized to a no-antibody control set to 100% (OD 1.2-1.5) and is presented as relative signal (%). Data are shown as mean ±SEM (n = 15) from 5 independent experiments. **(D)** Indicated pelleted rVSVs, prenormalized by their VSV M contents, were subjected to SDS-PAGE and silver-staining to visualize incorporated proteins. A representative image from 3 independent experiments is shown. Hantavirus Gn and Gc, as well as VSV N and M proteins, are indicated. The magenta arrowhead indicates the VSV G protein. **(E)** Representative immunoblot from 3 independent experiments showing virion-incorporated Gc proteins in pelleted rVSV-ANDV and rVSV Necoclí particles. Some error bars are smaller than the symbols or bar borders.

We next tested whether the EC1-specific mAb-3305 could block this interaction. Preincubation of sEC1-2-coated plates with mAb-3305 before addition of rVSVs caused a dose-dependent decrease in binding of all 3 rVSVs, like rVSV-ANDV, whereas an unrelated human IgG-Ctrl did not (Fig. 4C). For rVSV-Nova, binding to sEC1-2 was weak, and we did not observe reproducible inhibition by mAb-3305 in the capture ELISA; however, because the baseline signal was near the lower limit of detection, blockade could not be assessed reliably (data not shown).

Silver-staining of virion-associated proteins after SDS-PAGE revealed that the lower binding observed for these rVSVs was not due to reduced incorporation of Gn/Gc into particles (Fig. 4D). This was further supported by Western blotting of pelleted rVSV-Necoclí using antisera raised against an ANDV Gc peptide with 100% sequence identity (Fig. 4E). Collectively, these results indicate that Necoclí, Tula, and Nova Gn/Gc specifically bind PCDH1, albeit with different apparent avidities, with Nova showing the weakest binding.

### PCDH1 EC1 targeting blocks Necoclí and Tula Gn/Gc-mediated infection

PCDH1 binds New World hantavirus Gn/Gc via its EC1 domain, and soluble EC1 and EC1-2, as well as the EC1-specific mAb-3305, neutralize these infections(17, 18). To determine whether targeting the PCDH1:Gn/Gc interaction also blocks Necoclí, Tula, and Nova virus infection, we generated a human Fc-tagged version of PCDH1 EC1 (EC1-Fc), expressed it in ExpiCHO cells by transient transfection, and purified it from culture supernatants by affinity chromatography. Silver staining and immunoblotting after SDS-PAGE confirmed that EC1-Fc was highly purified (Fig. 5A). Preincubation of rVSVs with EC1-Fc for 1 hour at 37°C inhibited infection of HPMECs by rVSV-ANDV and rVSV-Necoclí with similar efficiency in a dose-dependent manner (Fig. 5B). Compared with rVSV-HTNV, the negative control, rVSV-Tula and rVSV-Nova were also neutralized, although more weakly, consistent with their lower apparent binding avidity for PCDH1 (Fig. 5B).

**Fig. 5.**
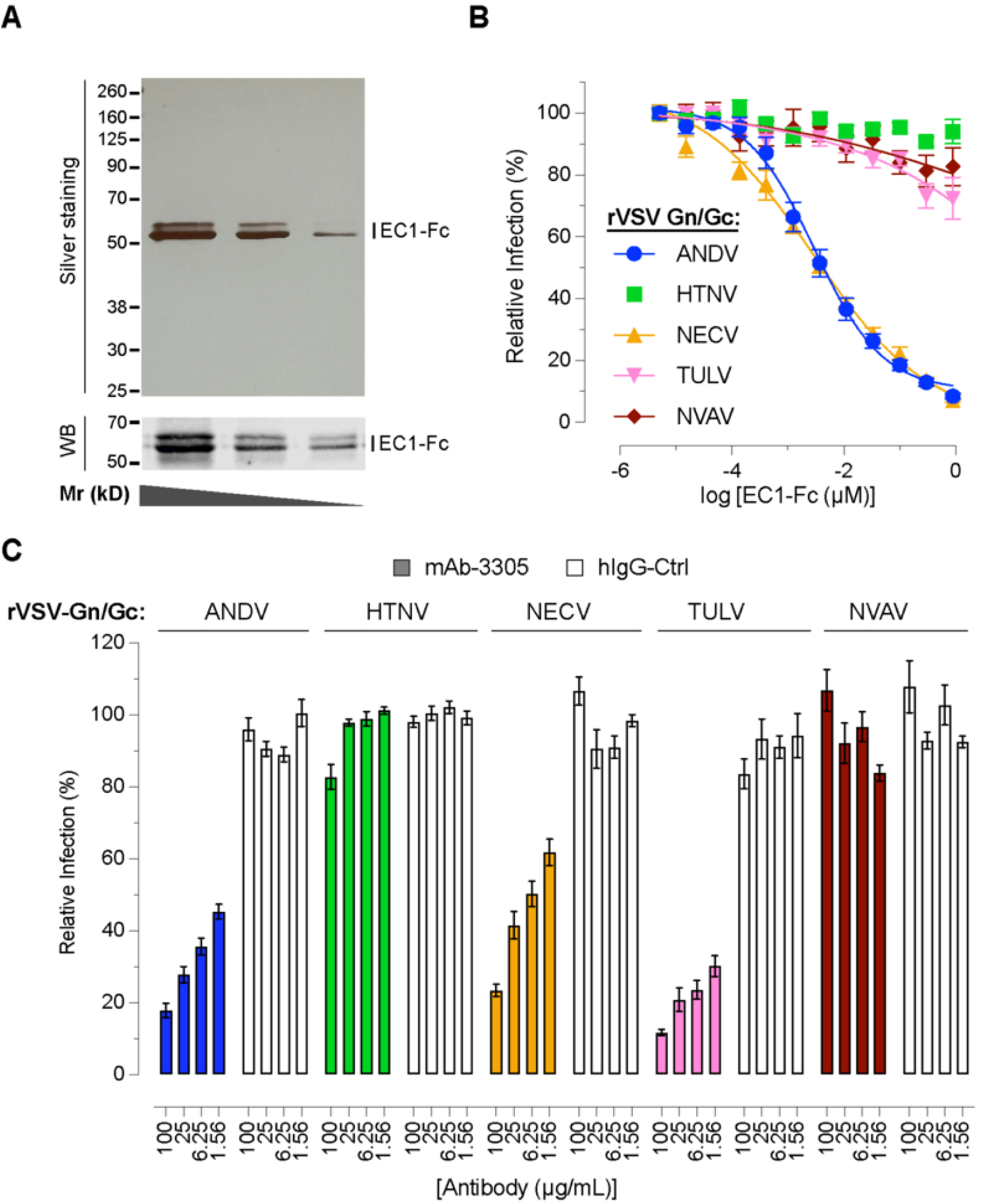
PCDH1 EC1 targeting blocks Necoclí and Tula Gn/Gc-mediated infection. **(A)** Affinity-purified EC1-Fc protein expressed in ExpiCHO cells by transient transfection was analyzed by SDS-PAGE followed by silver staining (top) or Western blotting (bottom) with an anti-human IgG antibody. Representative images from 3 independent experiments are shown. **(B)** Fixed amounts of the indicated rVSVs, pretitrated to achieve approximately 20% infection in untreated control wells, were preincubated with serial 3-fold dilutions of EC1-Fc (starting at 0.9 μM) for 1 hour at 37 °C before infection of HPMECs. Cells were fixed 16 hours later and infection was quantified as in Fig. 2C. Data are presented as relative infection, with no EC1-Fc set to 100%. Data are shown as mean ± SEM (n = 12-24) from 3-6 independent experiments. **(C)** HPMECs were preincubated for 1 hour at 37°C with the indicated concentrations of EC1-specific mAb-3305 or an unrelated hIgG-Ctrl and then infected with the indicated rVSVs, which were pretitrated to achieve approximately 20% infection in untreated control wells. Cells were fixed 16 hours later and infection was quantified as in Fig. 2C. Data are presented as relative infection, with no antibody set to 100%. Data are shown as mean ± SEM (n = 5-6) from 3 independent experiments). Each n in panels B and C represents the infection status of 500-1,000 cells per condition. Some error bars are smaller than the symbols or bar borders.

We next evaluated the neutralization activity of the EC1-specific mAb-3305 against these rVSVs by preincubating HPMEC with antibody or control IgG for 1 hour before infection, as in our previous work (Fig. 5C)(18). rVSV-ANDV showed dose-dependent inhibition by mAb-3305, whereas an unrelated hIgG-Ctrl did not. rVSV-HTNV showed no inhibition. rVSV-Necoclí and rVSV-Tula also showed dose-dependent inhibition, with rVSV-Tula being more efficiently neutralized by mAb-3305 (Fig. 5C). In contrast, rVSV-Nova was not neutralized by mAb-3305. Together, these data demonstrate that the Gn/Gc proteins of Necoclí and Tula engage with the EC1 domain of PCDH1 in a manner that can be blocked by both soluble EC1-Fc and the EC1-specific mAb-3305, whereas Nova shows only partial sensitivity to PCDH1-targeting approaches.

### Authentic Tula virus infection is PCDH1-dependent

The availability of hantavirus isolates and reagents for infection studies remains limited. Among the viruses included in our panel, we examined the effect of PCDH1 knockout on authentic Tula virus infection in TIME endothelial cells. WT TIME cells, PCDH1 knockout (KO) TIME cells carrying empty vector, and PCDH1 KO TIME cells complemented with PCDH1 cDNA were infected with Tula virus, fixed 48 hours later, and immunostained for nucleoprotein expression to assess infection (Fig. 6A). Compared with WT cells, Tula virus infection was reduced by approximately 80% in PCDH1 KO cells and was restored by PCDH1 complementation (Fig. 6B). These findings corroborate the results obtained with rVSV-Tula and show that authentic Tula virus infection is PCDH1-dependent.

**Fig. 6.**
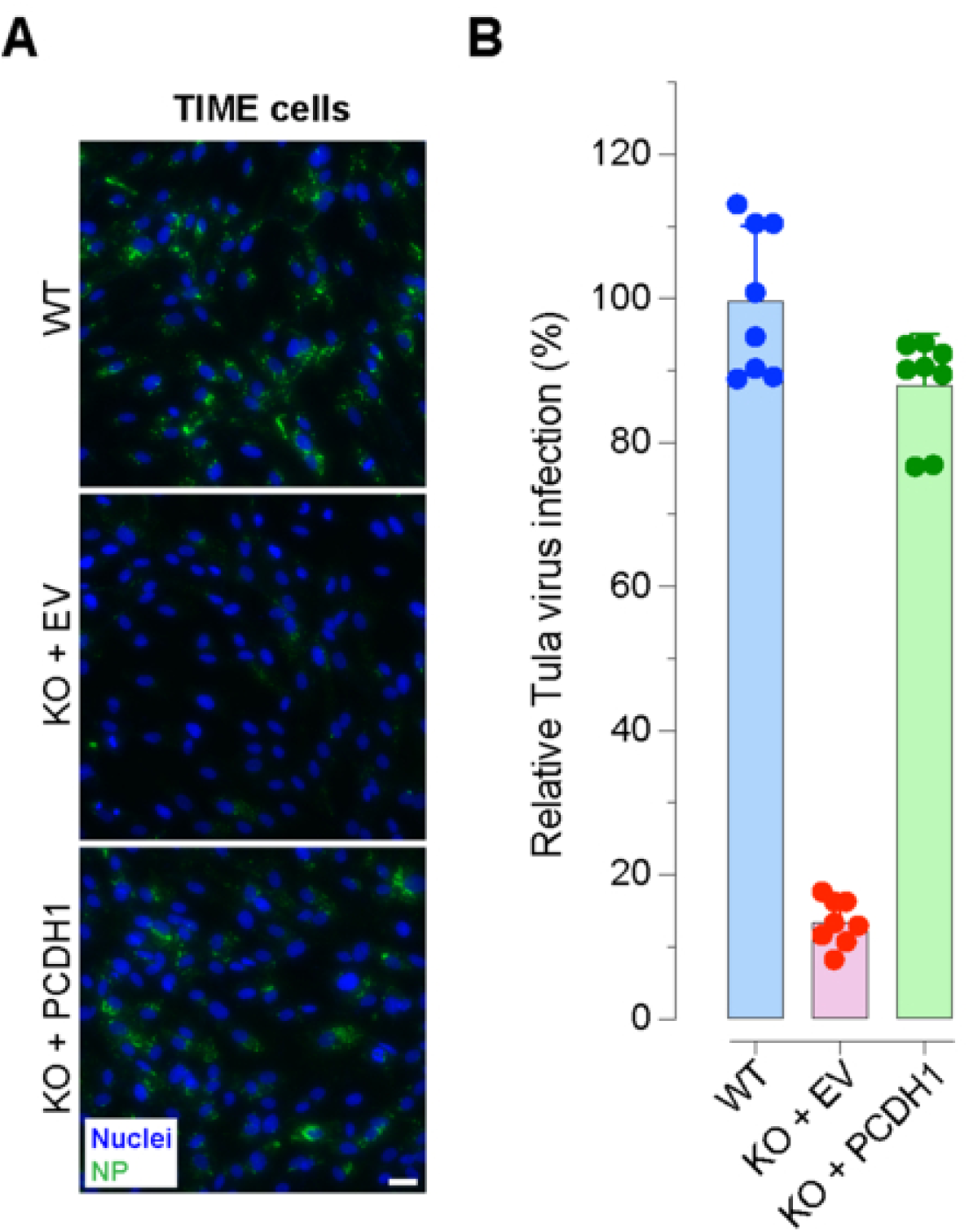
Authentic Tula virus infection is PCDH1-dependent. **(A)** WT TIME cells, a mixed PCDH1 KO TIME cell line carrying empty vector (KO + EV), and PCDH1 KO TIME cells complemented with PCDH1 cDNA (KO + PCDH1) were infected with authentic Tula virus and fixed 48 hours later. Cells were immunostained for nucleoprotein expression, and nuclei were counterstained with Hoechst 33342. Four images per well were acquired. Representative images from one of two independent experiments are shown. Scale bar = 30 μm. **(B)** Infected cells were quantified using Cytation-5 onboard Gen5 software, and data are presented as relative infection, with WT set to 100%. Each dot represents the infection measured from 200-300 cells per image. Data are shown as mean ± SEM (n = 8) from 2 independent experiments). Some error bars are smaller than the symbols.

### A broadly neutralizing human mAb targeting Gn/Gc (ADI-42898) binds and neutralizes these rVSVs

Because PCDH1-targeting approaches are ineffective against PCDH1-independent viruses, we tested the broadly neutralizing human anti-Gn/Gc ADI-42898 for activity against all seven rVSVs included in this study. Given the sequence heterogeneity of hantavirus Gn/Gc, broadly neutralizing antibodies are rare. However, ADI-42898 has shown potent activity against multiple New World and Old World hantaviruses *in vitro* and in animal models, despite recognizing a quaternary epitope located in a relatively variable region of Gn/Gc(41, 42).

An alignment of the mapped ADI-42898 epitope, encompassing the Gn-capping loop and the bc/cd loops of Gc, across hantaviruses revealed substantial variation at residues corresponding to Q98, E750, and E768 of PUUV Gn/Gc (Fig. 7A). We next evaluated binding of ADI-42898 to plate-captured rVSVs normalized for particle input by VSV matrix (M) protein levels and observed marked differences among the rVSVs (Fig. 7B). Compared with rVSV-ANDV and rVSV-HTNV, binding to rVSV-Kenkeme, rVSV-Necoclí, and rVSV-Nova was weaker, in that order. Binding to rVSV-Sangassou was low but was reproducibly above the positive cutoff defined in Fig. 7B. In contrast, binding to rVSV-Oxbow, rVSV-Tula, and rVSV-Thottapalayam was below the limit of detection (Fig. 7B). Finally, we evaluated the neutralization activity of ADI-42898 and the negative control hIgG-Ctrl against all seven rVSVs, together with positive controls (rVSV-ANDV and rVSV-HTNV) and the negative control virus VSV-G, in Vero cells. The hIgG-Ctrl did not neutralize any of these viruses (Fig. 7C). Consistent with the binding data, ADI-42898 efficiently neutralized rVSV-Kenkeme, rVSV-Necoclí, rVSV-Nova, and rVSV-Sangassou (Fig. 7C). Neutralization of rVSV-Oxbow, rVSV-Tula, and rVSV-Thottapalayam was weak or undetectable. In sum, these data show that ADI-42898 retains neutralizing activity against several highly divergent hantavirus Gn/Gc proteins and that this rVSV panel provides a useful system for evaluating entry inhibitors and mechanisms of cellular entry.

**Fig. 7.**
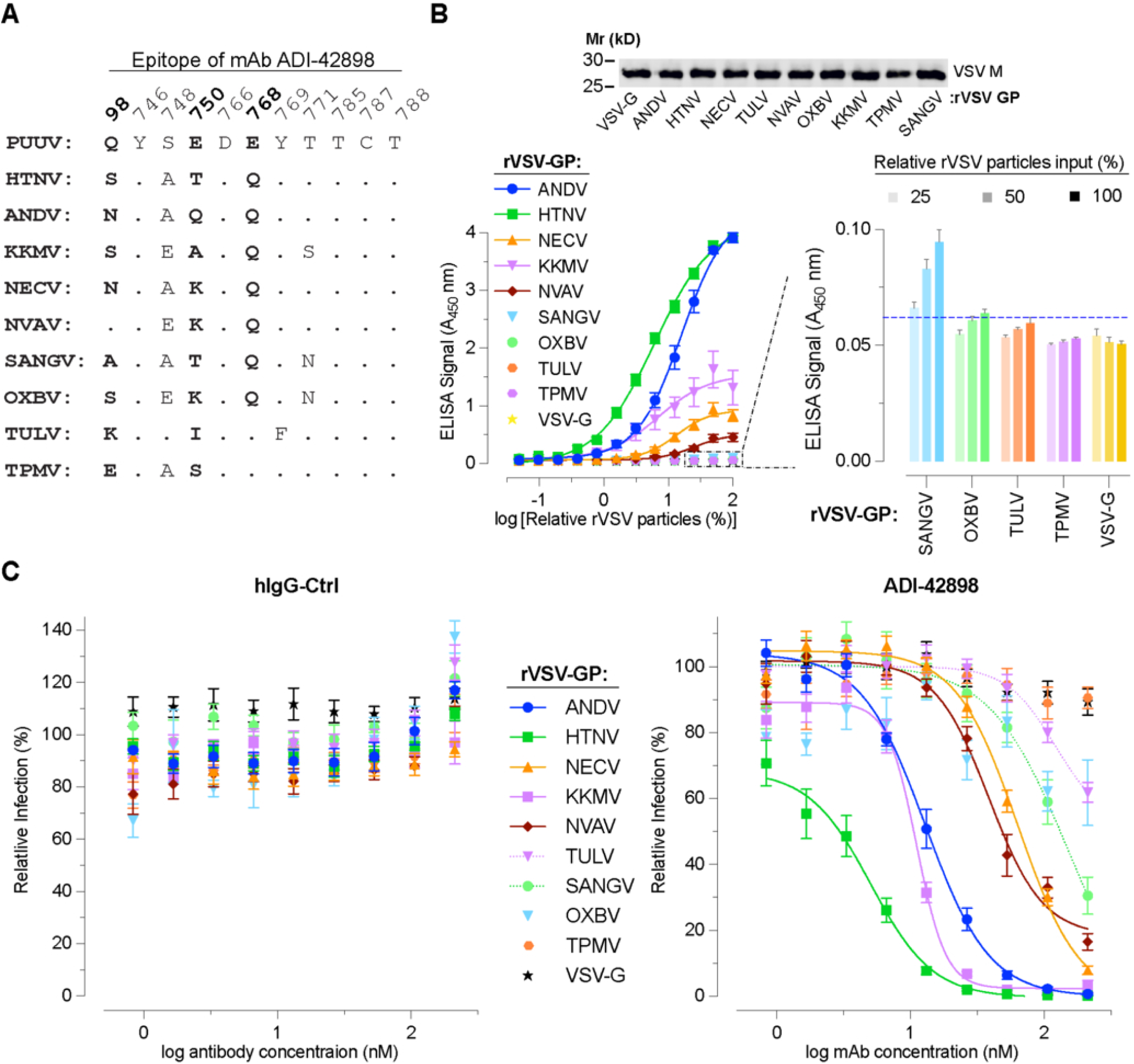
A broadly neutralizing human mAb targeting Gn/Gc (ADI-42898) binds and neutralizes these rVSV. **(A)** Alignment of the amino acid sequences corresponding to the mapped ADI-42898 epitope across hantavirus Gn/Gc, including Puumala virus (PUUV), Andes virus, and Hantaan virus, generated using SnapGene v8.2.2. Key functional PUUV residues Q98, E750, and E768 of PUUV within the epitope are shown in bold. Amino acid changes in other Gn/Gc are indicated. Dots indicate amino acids identical to those of PUUV. **(B)** Top, representative Western blot showing normalized amounts of the indicated pelleted rVSV particles, estimated by VSV matrix (M) protein levels, used as input for the ADI-42898 ELISA. Bottom, plates coated with serial twofold dilutions of prenormalized rVSVs were incubated with the ADI-42898 at 1 μg/mL and bound antibody was quantified by ELISA using an HRP-conjugated anti-human secondary antibody. Right lower panel, the same data are shown an a bar graph to illustrate weak but reproducible binding to rVSV-Sangassou more clearly. Data are shown as mean ± SEM (n = 9-12) from 3-4 independent experiments. **(C, D)** Fixed amounts of the indicated rVSVs, pretitrated to achieve 20-30% infection in untreated control wells, were pre-incubated with serial twofold dilutions of an unrelated hIgG-Ctrl (C) or ADI-42898 (D), starting at 30 nM for 1 hour at 37 °C before infection of African green monkey kidney Vero cells. Cells were fixed 8 hours later and infection was quantified as in Fig. 2C. Data are presented as relative infection, with no antibody set to 100%. Data are shown as mean ± SEM (n = 9-12) from 3-4 independent experiments. Some error bars are smaller than the symbols.

## Discussion

Hantavirus disease is fundamentally an endothelial disease, with vascular leakage rather than overt cytolysis driving much of the clinical pathology, making endothelial entry a biologically relevant phenotype for comparative study(43, 44). Replication-competent rVSV systems therefore provide a practical and informative way to isolate glycoprotein-mediated entry under lower biocontainment. We previously established that the HTNV adaptive substitutions I532K and S1094L enhance rVSV infectivity by promoting Gn/Gc localization to the plasma membrane and increasing glycoprotein incorporation into virions(16). Earlier VSV pseudotype systems bearing HTNV or SEOV glycoproteins were likewise developed as rapid, safer surrogate assays, underscoring the broader utility of VSV-based platforms for entry and antiviral studies(45, 46). In this context, our data show that Gn/Gc proteins from multiple divergent hantaviruses can mediate infection of primary human endothelial cells (Fig. 2), indicating that endothelial cell competence is broader than generally appreciated. This interpretation is also consistent with a recent large-scale survey of animal virus entry(47).

Our results further indicate that endothelial entry competence is not binary, but highly graded. Most of the rVSVs in our panel infected HPMECs, yet did so with markedly different efficiencies and temporal profiles, suggesting that many hantavirus glycoproteins can access human entry pathways but vary substantially in how effectively they do so (Fig. 2). Because Gn/Gc is the only hantavirus determinant in this system, these phenotypic differences most likely reflect variation in receptor engagement, uptake, fusion efficiency, or glycoprotein-driven spread after initial infection. The capacity to enter human endothelial cells should not be equated with zoonotic potential or pathogenicity because disease outcomes also depend on post-entry replication, innate immune interactions, endothelial dysfunction, and host ecology, all of which lie beyond the scope of an entry-only platform(43, 44).

A key finding is that several divergent Gn/Gc proteins functioned in the rVSV platform without acquiring additional rescue-associated mutations, despite the evolutionary distance among the viruses studied (Figs. 1 and 2). This is conceptually important because it suggests that, once properly expressed and incorporated, many hantavirus glycoproteins are intrinsically compatible with mediating infection in human cells in the VSV background. Because several constructs (Oxbow, Sangassou, Kenkeme, Thottapalayam, and Nova) were pre-engineered with HTNV-like rescue-enhancing substitutions, our data do not show that all wild-type glycoproteins would behave identically in the absence of adaptation. Rather, they show that extensive additional adaptation is generally not required beyond that starting point. In that sense, the study strengthens the view that cellular entry by hantavirus Gn/Gc is comparatively permissive, while still leaving open the possibility that later steps in infection impose stronger species barriers.

The principal mechanistic advance of this study is the expansion of PCDH1 usage beyond the canonical model centered on New World hantaviruses. PCDH1 was identified as an essential entry receptor for ANDV and SNV in pulmonary endothelial cells, and that work showed that hantavirus glycoproteins directly recognize the outermost EC1 domain of PCDH1. Importantly, PCDH1 ablation also protected hamsters from lethal ANDV challenge(18). Subsequent work mapped the critical receptor-contact surface more precisely and showed that two-point mutations in PCDH1 disrupt hantavirus recognition and protect against lethal infection, firmly establishing PCDH1 as a bona fide entry receptor rather than a passive attachment factor(17). In parallel, CRISPR-based genetic depletion studies in human endothelial cells found that loss of PCDH1, but not several previously proposed receptors, substantially reduced entry and infection by a subset of virulent hantaviruses, reinforcing the importance of PCDH1 in endothelial cells specifically(25). Our identification of Necoclí, Tula, and Nova virus Gn/Gc here as functionally PCDH1-dependent broadens the phylogenetic distribution of hantaviruses that can use this receptor and argues against a simple Old World/New World receptor dichotomy.

Equally important, our data suggest that PCDH1 usage phenotypes were not uniform across the viruses tested. Necoclí virus most closely resembled the prototypic PCDH1-dependent phenotype established for ANDV (Fig. 3), whereas Tula virus and especially Nova virus showed weaker or more selective behavior across binding and inhibition assays (Figs. 4 and 5). That pattern argues that these viruses are not simply weaker versions of the same receptor interaction. Instead, the combined knockout, complementation, binding, and neutralization results are more consistent with distinct modes of EC1 engagement that differ in avidity, geometry, or epitope accessibility. The Nova virus phenotype is particularly informative: detectable PCDH1 dependence and sEC1-2 binding were accompanied by resistance to mAb-3305 neutralization and the inability to assess mAb-3305 blockade reliably in the capture ELISA because of weak baseline binding. This suggests productive engagement of PCDH1 that is nevertheless not equally disrupted by all EC1-targeting reagents. A conceptually similar pattern emerged in our recent study of bat cells, in which rVSV-ANDV entry in *Carollia perspicillata* kidney cells was inhibited by sEC1-2 but not by mAb-3305(48). An alternative interpretation is that Nova virus relies only partly on PCDH1, with non-PCDH1 receptor(s) or entry-promoting cofactors contributing more prominently to infection. This interpretation is also consistent with prior receptor-mapping studies showing that hantavirus recognition depends on a defined surface on PCDH1 EC1 and that changes within this surface can influence glycoprotein recognition and cell entry(17).

The discrepancy between U2OS and TIME complementation data reinforces the view that receptor usage is cell-context-dependent (Fig. 3). In clonal U2OS cells, complementation incompletely restored some phenotypes, whereas in TIME endothelial cells, rescue was more evident (Fig. 3). An alternative explanation is that the incomplete rescue seen in U2OS cells reflects a clone-specific phenotype rather than a general feature of PCDH1 dependence, which would make the TIME endothelial-cell data more informative result. The most parsimonious interpretation is therefore that receptor abundance, trafficking, glycosylation, membrane organization, or additional endothelial cofactors modulate how efficiently a given Gn/Gc exploits PCDH1. This interpretation is consistent with prior studies that identified PCDH1 as a hantavirus receptor in pulmonary endothelial cells, supporting the idea that receptor usage is best evaluated in cell types relevant to hantavirus tropism(18, 25). The authentic Tula virus result is therefore especially valuable (Fig. 6) because it extends the central rVSV-based conclusion to authentic virus infection. These findings also have immediate implications for countermeasure design. Since only a subset of viruses in this panel are PCDH1 dependent, host-directed strategies aimed at blocking PCDH1 are unlikely to be pan-hantaviral. Even among PCDH1-using viruses, the differing responses to soluble EC1-Fc and mAb-3305 (Fig. 5) suggest that not all receptor-targeted modalities will perform equivalently. Residual infection after PCDH1 loss is also consistent with prior studies, which established PCDH1 as a bona fide hantavirus entry receptor while also indicating PCDH1 ablation or targeting does not invariably eliminate infection in every experimental setting(17, 18, 25). Collectively, these findings support a model in which PCDH1 is an important determinant of susceptibility, but not necessarily the sole determinant, and they argue that additional attachment factors or entry-promoting cofactors remains to be identified, especially for viruses that appear PCDH1-independent.

The ADI-42898 data highlight both the breadth and the limits of broad anti-Gn/Gc neutralization (Fig. 7). ADI-42898 was previously shown to recognize a quaternary epitope on the Gn/Gc complex, block entry of seven HCPS- and HFRS-associated hantaviruses, and protect animals against both ANDV and PUUV challenge, making it one of the strongest reported pan-hantavirus antibody leads(41). Structural follow-up work explained this breadth by showing that the antibody bridges the Gn/Gc heterodimer, engages conserved Gc fusion-loop features together with the Gn capping region, and stabilizes the complex in its prefusion state. It also showed that reduced potency against divergent viruses can arise from altered binding behavior under endosomal acidic pH(42). In that context, our observation that ADI-42898 neutralizes several, but not all, of these newly generated rVSVs is biologically informative. It shows that broad cross-clade activity is achievable, but also that epitope presentation and structural divergence still impose meaningful limits on breadth. Taken together, our studies support a model in which human endothelial cell susceptibility to hantavirus Gn/Gc-mediated infection is broader than previously assumed, PCDH1 usage is more widely distributed but not universal, and broad neutralization across phylogenetic distant hantaviruses is achievable yet incomplete.

## Materials and Methods

### Cells

Human embryonic kidney 293FT and human hepatoma Huh7.5.1 cells were maintained in high-glucose Dulbecco’s modified Eagle medium (DMEM), and U2OS cells were maintained in McCoy’s 5A medium. Both media were supplemented with 10% fetal bovine serum (FBS), 1% GlutaMAX, and 1% penicillin-streptomycin. Telomerase-immortalized human microvascular endothelial (TIME) cells were recovered in Lonza EBM-2 medium (EBM™-2 Basal medium supplemented with EGM™-2 SingleQuots™ Supplements) and maintained in Lifeline VascuLife medium (Lifeline VascuLife® Basal medium supplemented with VascuLife® VEGF LifeFactors Kit). Primary human pulmonary microvascular endothelial cells (HPMECs) were recovered and maintained in the previously mentioned VascuLife medium. African green monkey kidney Vero cells were cultured in high-glucose DMEM supplemented with 2% FBS, 1% GlutaMAX, and 1% penicillin-streptomycin.

For expression of soluble proteins and antibodies, Freestyle 293-F (293FS) and Chinese hamster ovary ExpiCHO-S cells were maintained in non-baffled 250-ml shaker flasks. 293FS cells were cultured in Gibco FreeStyle 293 Expression medium supplemented with 1% penicillin-streptomycin and 1% GlutaMAX. ExpiCHO-S cells were cultured in Gibco’s ExpiCHO Expression medium supplemented with 1% penicillin-streptomycin and 1% GlutaMAX.

Adherent cell lines were maintained at 37°C in a humidified incubator with 5% CO_2_. Suspension 293FS and ExpiCHO-S cells were maintained at 37°C in a humidified incubator with 8% CO_2_ on an orbital shaker at 125 to 130 rpm. Cells were tested for mycoplasma every month and were negative.

The generation and characterization of PCDH1 knockout (KO) U2OS and TIME cells, as well as PCDH1 KO U2OS and TIME cells complemented with PCDH1 cDNA were described previously(18, 25). PCDH1-complemented KO U2OS cells were maintained in medium supplemented with puromycin (2 μg/mL).

### Generation of rVSV Gn/Gc

Replication-competent recombinant vesicular stomatitis viruses (rVSVs) expressing enhanced green fluorescent protein (eGFP) and bearing hantavirus glycoproteins were generated as described previously(18, 49–51). Briefly, glycoprotein precursor sequences of Necoclí virus (NECV; GenBank accession no. KF494345.1), Tula virus (TULV; Z69993.1), Nova virus (NVAV; KR072622.1), Oxbow virus (OXBV; FJ539167.1), Kenkeme virus (KKMV; KJ857337.1), Thottapalayam virus (TPMV; DQ825771.1), and Sangassou virus (SANGV; JQ082301.1) were codon optimized for expression in human cells and synthesized by Twist Biosciences. These sequences were cloned between MluI and NotI restriction sites to replace the native VSV glycoprotein G gene in the plasmid encoding the cognate VSV genome under the control of the T7 polymerase promoter. After sequence verification, the genome plasmids, together with the helper plasmids encoding VSV N, P, M, G and L and T7 RNA polymerase, were transfected into 293FT cells using polyethylenimine (PEI). Supernatants were then transferred to Huh7.5.1 cells for amplification and generation of rVSV stocks. Every virus stock was validated by sequencing the entire Gn/Gc coding region following genomic RNA isolation and RT-PCR. The generation of rVSV-ANDV and rVSV-HTNV was described previously(49).

For ELISA, Western blotting and silver staining experiments, rVSV particles in clarified supernatants were pelleted by ultracentrifugation for 2 hours at 20,000 rpm and 4°C in an SW32 rotor. Pellets were resuspended in 1x NT buffer overnight at 4°C, layered over a 10% sucrose cushion, and centrifuged again for 2 hours at 20,000 rpm and 4°C in an SW41 rotor. The resulting pellets were resuspended in 1x NT buffer, aliquoted, and stored at -80°C.

### Authentic Tula virus infection

To examine authentic virus infection, WT TIME cells, PCDH1 KO TIME cells carrying empty vector, and PCDH1 KO cells complemented with PCDH1 cDNA were infected with Tula virus (TULV) at a multiplicity of infection of 0.1. At 48 hours post-infection, cells were fixed with 3% paraformaldehyde and immunostained for nucleoprotein expression using a rabbit polyclonal anti-Puumala virus nucleocapsid serum(52), followed by Alexa Fluor-488-conjugated donkey anti-rabbit IgG. Four images were acquired per well on a Cytation-5 imager, and infected cells were quantified using Gen5 software (Agilent BioTek). Relative infection was normalized to WT cells, which were set to 100%.

### Generation of PCDH1 EC1-Fc plasmid

To generate a human Fc-tagged version of PCDH1 EC1 (EC1-Fc), a codon-optimized sequence encoding amino acids 58 to 165 of human PCDH1 (GenBank accession number NM_002587) was cloned in frame downstream of the signal peptide sequence MGWSCIILFLVATATGAHS in the pMAZ-IgH vector encoding a human Fc region using NEBuilder HiFi DNA assembly. The resulting plasmid, pMAZ-PCDH1-EC1-Fc, was sequence verified by nanopore sequencing (Plasmidsaurus).

### Expression of soluble proteins and antibodies

For expression of EC1-Fc, mAb-3305, and ADI-42898, ExpiCHO-S cells were used. One day before transfection, cells were split to 3 × 10^6^ to 4 × 10^6^ cells/mL. On the day of transfection, cells were adjusted to 6 × 10^6^ cells/mL and transfected with 0.8 μg/mL of plasmid DNA using ExpiFectamine CHO reagent in OptiPRO SFM; a single plasmid was used for EC1-Fc, whereas equal amount of heavy- and light-chain plasmids for mAb-3305 and ADI-42898. At 18 to 22 hours post-transfection, ExpiFectamine CHO enhancer and ExpiCHO feed were added. Cells were then returned to 37°C, 8% CO_2_, and continued to shake at 125 to 130 rpm. Secreted proteins and antibodies were harvested 7 to 10 days post-transfection(18).

Soluble PCDH1 EC1-2 (sEC1-2) was expressed in 293FS cells as described previously(18). Briefly, cells were seeded at 5 × 10^5^ to 6 × 10^5^ cells/mL one day before transfection and adjusted to 1 × 10^6^ viable cells/mL on the day of transfection. Cells were transfected with 0.67 μg/mL of the sEC1-2 expression plasmid using PEI. The secreted protein was harvested 6 days later.

### Purification of EC1-Fc and antibodies

For purification of EC1-Fc, mAb-3305 and ADI-42898, ExpiCHO-S cultures were centrifuged at 4,000 rpm for 20 minutes at 4°C and supernatants were adjusted to pH 8.0 with NaOH. Complete Mini Protease Inhibitor (Thermo Scientific) was added and clarified supernatants were incubated overnight at 4°C with 0.5 mL packed Pierce Protein A agarose beads pre-equilibrated in Gentle Ag/Ab binding buffer (Thermo Scientific). The following day, beads were loaded onto Econo-Pac chromatography columns (Bio-Rad), washed with Pierce Gentle Ag/Ab Binding Buffer, and proteins were eluted with Pierce Elution Buffer (Thermo Scientific). Eluates were buffer exchanged into HEPES-NaCl buffer, pH 7.4, using PD-10 desalting columns and concentrated with 30-kDa-molecular-weight-cutoff Amicon concentrators. Protein concentrations were measured by absorbance at 280 nm. Aliquots of purified proteins were stored at -80°C.

### Purification of soluble PCDH1 EC1-2

For purification of sEC1-2, 293FS cultures were centrifuged at 1,500 rpm for 15 minutes at 4°C and clarified supernatants were incubated overnight at 4°C with 1.5 mL of packed Ni-NTA Agarose beads (Qiagen), and Complete Mini Protease Inhibitor. Beads were recovered on Econo-Pac chromatography columns (Bio-Rad), washed with PBS and then with wash buffer containing 20 mM imidazole, 50 mM NaH_2_PO_4_ (pH 8.0) and 300 mM NaCl, and proteins were eluted with buffer containing 250 mM imidazole, 50 mM NaH_2_PO_4_ (pH 8.0) and 300 mM NaCl. Eluted proteins were buffer exchanged into PBS and concentrated with 10-kDa-molecular-weight-cutoff Amicon concentrators. Protein concentrations were measured by absorbance at 280 nm. Aliquots of purified protein were stored at -80°C. Purified proteins and antibodies were analyzed by SDS-PAGE followed by silver staining (for purity) and Western blotting (for identity), ELISA or neutralization assays, as appropriate.

### sEC1-2 capture and competition ELISAs

These ELISAs were performed essentially as described previously(17, 18). Briefly, purified, pelleted rVSVs were normalized by VSV matrix (M) protein levels. High-binding 96-well ELISA plates (Corning) were coated overnight at 4°C with sEC1-2 (100 ng/well). Plates were blocked with 5% nonfat dry milk in 1x PBS for 1 hour at 37 °C. rVSV particles were labeled with the short-chain phospholipid probe FSL-biotin (Sigma-Aldrich) for 1 hour at 37 °C. The labeled rVSVs were serially diluted twofold in 5% milk in PBS and added to the plate for 1 hour at 37 °C. Plates were washed three times with PBS, incubated with streptavidin-HRP in 5% milk in PBS for 1 hour at 37°C, washed again, and developed with 1-Step™ TMB substrate (Thermo Scientific). Reactions were stopped with 0.5 M H_2_SO_4_, and absorbance was measured at 450 nm using a Cytation-5 plate reader (Agilent BioTek). ELISA data were fit in GraphPad Prism v11 to a four-parameter logistic (4PL) sigmoidal/variable-slope model with log-transformed concentration on the x-axis. For competition ELISAs, sEC1-2-coated plates were incubated with the indicated fivefold dilutions of mAb-3305 or a nonspecific human IgG-Ctrl antibody before addition of rVSVs. In these experiments, a fixed normalized amount of rVSV was used instead of serial dilutions.

### ADI-42898 binding ELISA

ADI-42898 binding ELISAs were performed as described previously with minor modifications(41). Briefly, normalized purified rVSVs were serially diluted twofold in PBS and used to coat high-binding 96-well ELISA plates overnight at 4°C. Plates were blocked with 5% milk in PBS for 1 hour at 37°C and then incubated with ADI-42898 at 1 μg/mL for 1 hour at 37°C. After washing, bound antibody was detected with HRP-conjugated anti-human IgG secondary antibody in 5% milk in PBS, and signal was developed and measured as described above. ELISA data were fit in GraphPad Prism v11 to a four-parameter logistic (4PL) sigmoidal/variable-slope model with log-transformed concentration on the x-axis.

### Neutralization assays

For mAb-3305 neutralization assays were performed as described previously(18). HPMECs were seeded in a 96-well plate at approximately 8,000 cells per well 1 day before infection. Cells were preincubated for 1 hour with the indicated concentrations of mAb-3305 or human IgG-Ctrl and then infected with pretitrated amounts of rVSV yielding approximately 10 to 30% baseline infection. At 8 hours post-infection, cells were fixed with 3% formaldehyde, and nuclei were stained with Hoechst 33342. Infection was quantified by automated counting of eGFP-positive cells as described above.

For EC1-Fc neutralization assays were described previously(18). Briefly, pretitrated rVSVs were incubated for 1 hour at 37°C with the indicated concentrations of EC1-Fc (starting concentration, 0.9 μM; threefold serial dilutions) before addition to HPMECs. Cells were fixed 8 hours later, and infection was quantified as described above.

For ADI-42898 neutralization assays were performed as described(41, 42). Vero cells were seeded in a 96-well plate at approximately 8,000 cells per well 1 day before infection. Pretitrated rVSVs yielding approximately 20-40% baseline infection were incubated for 1 hour at 37°C with the indicated concentrations of ADI-42898 or human IgG-Ctrl (starting concentration, 30 nM; twofold serial dilutions). Virus-antibody mixtures were then added to Vero cells for 8 hours and infection was quantified as described above. The EC1-Fc and ADI-42898 neutralization data were fit in GraphPad Prism v11 to a four-parameter (4PL) sigmoidal/variable-slope model using log-transformed concentration on the x-axis.

### Western blotting

Pelleted rVSV particles normalized by VSV-M content were denatured and resolved on 10% SDS-PAGE gels. Proteins were transferred to nitrocellulose membranes, which were blocked in 5% milk in PBS-T. Membranes were incubated overnight at 4°C with mouse anti-VSV-M antibody (clone 23H12) or rabbit anti-Gc serum raised against the ANDV peptide EKDYQYETGWGC (Pacific Immunology; 1:100). Membranes were washed and incubated for 1 hour at room temperature with IRDye-conjugated goat anti-mouse or goat anti-rabbit secondary antibodies (LI-COR; 1:2,000). Proteins were visualized using an Odyssey CLx Infrared Imaging System (LI-COR).

### Silver staining

Pelleted rVSV particles normalized by VSV-M content were resolved on 10% SDS-PAGE gels and subjected to silver staining using a modified carbonate silver stain based on the Blum method(53). Gels were fixed overnight at 4°C in a fixative containing 50% methanol, 12% acetic acid, 0.0175% formaldehyde and Milli-Q water, washed in 35% ethanol, and sensitized with 0.02% sodium thiosulfate. Gels were then rinsed with Milli-Q water, incubated in 0.2% silver nitrate containing 0.0266% formaldehyde for 20 minutes at room temperature, briefly washed, and developed in 6% sodium carbonate, 0.0175% formaldehyde developer containing 0.0004% sodium thiosulfate until bands became visible. Development was stopped in 50% methanol/12% acetic acid, and gels were visualized using an I2R Glow Box transilluminator.

### Statistics and software

Statistical details, including sample size, error bars, statistical tests, and exact p values are provided in the figures and figure legends. Statistical analysis was performed using GraphPad Prism v11. SnapGene v8.2.2 was used for plasmid map visualization and sequence analysis for molecular cloning, and Geneious Prime v2025.2 was used to generate the neighbor-joining phylogenetic tree.

## Data Availability

No new nucleotide or amino acid sequences, structural data or custom code were generated in this study. All data supporting the findings of this study are available within the article. Reagents generated in this study will be made available from the corresponding author upon reasonable request, subject to institutional and biosafety requirements.

## Acknowledgments

This work was supported by the National Institutes of Health (R21AI156482 to R.K.J., R01AI132633 to K.C., R56AI175292 to K.C. and R.K.J.). This work was also supported in part by a National Institute of General Medical Sciences (NIGMS) P20GM134974 grant awarded to the Center for Applied Immunology and Pathological Processes (CAIPP). We acknowledge the support of Research Core Facility Genomics Core (RRID: SCR_024775), Sushma Bharrhan in the CAIPP Immunophenotyping Core (RRID: SCR_024781), and resources from the CAIPP Bioinformatics and Modeling Core (RRID: SCR_024779). We would like to thank Tomas Strandin for providing the authentic Tula virus isolate and rabbit polyclonal anti-Puumala virus nucleocapsid serum. K.C. holds shares in Integrum Scientific, LLC and Eitr Biologics, Inc. K.C. and R.K.J. are co-inventors on a U.S. patent describing methods and assays for treating hantavirus infections.

